# Use of latent class analysis to identify multimorbidity patterns and associated factors in Korean adults aged 50 years and older

**DOI:** 10.1101/613646

**Authors:** Bomi Park, Hye Ah Lee, Hyesook Park

## Abstract

**Introduction:** Multimorbidity associated with significant disease and economic burdens is common among the aged. We identified chronic disease multimorbidity patterns in Koreans 50 years of age or older, and explored whether such patterns were associated with particular sociodemographic factors and health-related quality-of-life.

**Methods:** The multimorbidity patterns of 10 chronic diseases (hypertension, dyslipidemia, stroke, osteoarthritis, tuberculosis, asthma, allergic rhinitis, depression, diabetes mellitus, and thyroid disease) were identified via latent class analysis of data on 8,370 Korean adults aged 50+ years who participated in the sixth Korean National Health and Nutrition Examination Survey (2013-2015). The associations between multimorbidity patterns, and sociodemographic factors and health-related quality of life, were subjected to regression analysis.

**Results:** Three patterns of multimorbidity were identified: 1) a relatively healthy group (60.4% of the population); 2) a ‘cardiometabolic conditions’ group (27.8%); and, 3) an ‘arthritis, asthma, allergic rhinitis, depression, and thyroid disease’ group (11.8%). The female (compared to male) gender was associated with an increased likelihood of membership of the *cardiometabolic conditions* group (odds ratio [OR]=1.32, 95% confidence interval [CI]=1.15-1.51) and (to a much greater extent) the *arthritis, asthma, allergy, depression, and thyroid disease* group (OR=4.32, 95% CI=3.30-5.66). Low socioeconomic status was associated with membership of the two multimorbidity classes. Membership of the *arthritis, asthma, allergy, depression, and thyroid disease* group was associated with a significantly poorer health-related quality-of-life than was membership of the other two groups.

**Conclusion:** The co-occurrence of chronic diseases was not attributable to chance. Multimorbidity patterns were associated with sociodemographic factors and quality-of-life. Our results suggest that targeted, integrated public health and clinical strategies dealing with chronic diseases should be based on an understanding of multimorbidity patterns; this would improve the quality-of-life of vulnerable multimorbid adults.

## Introduction

Given aging populations, advances in medical care and public health policies, and improved living conditions, the co-occurrence of two or more chronic diseases in the same individual (multimorbidity) [1,2] is increasingly common [3-5]. A recent study of Koreans adults aged 50 and older found that more than one in four were multimorbid [6]. Multimorbidity is a public health concern, being associated with higher mortality, impaired functional status, a reduced quality-of-life, increased healthcare utilization, and a greater treatment burden [7-14]. The impact of multimorbidity can be much more complex (being synergistic) than the impact of individual diseases; health outcomes may differ by the disease combinations in play [3,15,16]. Thus, current single-disease-oriented public health strategies and clinical healthcare guidelines may be both incomplete and ineffective for multimorbid patients [17].

Certain chronic diseases tend to co-occur more often than expected by chance because they share pathophysiological pathways [18,19]. Identification of patterns of disease combinations and the characteristics of individuals exhibiting similar multimorbidity patterns may provide important information for policymakers and clinicians who seek to integrate strategic public health policy plans and healthcare management to address multimorbidity in at-risk groups more effectively. Even though multimorbidity has attracted increasing attention (because it is becoming the norm in the elderly; [17,20-25]), we still do not know how multiple chronic diseases cluster, the associated socioeconomic factors, and how multimorbidity affects quality-of-life.

Multimorbidity is highly complex. The use of a statistical approach to group a population into a limited number of subgroups with similar combinations of chronic diseases is much more practical than analysis of every possible disease combination. Therefore, we performed latent class analysis (LCA) based on the hypothesis that certain chronic diseases cluster. LCA identifies probabilistic rather than deterministic subgroups based on responses to a set of observed variables, and assumes that the pattern is explained by unobserved categorical latent variables of K classes [26,27]. Our objectives were: 1) to identify multimorbidity patterns in the general Korean population aged over 50 years using nationally representative survey data; and, 2) to explore whether such patterns were associated with certain sociodemographic characteristics and quality-of-life.

## Methods

### Data source and sample

We used data on adults 50 years of age and older who participated in the sixth Korean National Health and Nutrition Examination Survey (KNHANES) conducted in 2013-2015. KNHANES is a cross-sectional, nationally representative survey of the noninstitutionalized Korean population conducted by the Division of Chronic Disease Surveillance, Korea Centers for Disease Control and Prevention (KCDCP). KNHANES uses a stratified, multistage cluster sampling method based on geographical area, gender, and age. The details have been described previously [28].

### Measures

Chronic disease was based on self-reports, on whether participants had ever been physician-diagnosed with any disease on a pre-specified list. Analysis was limited to the 10 most common chronic diseases (hypertension, dyslipidemia, stroke, osteoarthritis, tuberculosis, asthma, allergic rhinitis, depression, diabetes mellitus, and thyroid disease; the prevalence of each is greater than 3% [29-31]) of the 28 diseases listed in the KNHANES. Multimorbidity was defined as the presence of two or more of these diseases in the same subject.

The sociodemographic variables evaluated included age, gender, household income, educational level, and occupation. The household income was the monthly income divided by the square root of household size [32], and was grouped into high and low using the median income as the cut-off. Educational level was categorized as low for those younger than 70 years who were high school graduates or less accomplished, and for those older than 70 years who were middle school graduates or less accomplished. In terms of occupational status, students and housewives were defined as unemployed; service workers. retailers, agriculture or fishery employees, technicians, mechanics, assemblers, and simple laborers were defined as manual workers; and managers, professionals, and office workers were considered to be non-manual workers. The health-related quality of life was assessed using the EuroQol 5 Dimensions (EQ-5D) instrument; this is a generic measure of health status. The EQ-5D features five dimensions (mobility, self-care, engagement in usual activities, pain/discomfort, and anxiety/depression); each dimension has three response options (1 = no problem, 2 = some problems, and 3 = a severe problem). The health-related quality of life was scored as a single value (the EQ-5D index score) using a validated algorithm [33,34].

### Statistical analysis

Disease prevalence, disease co-occurrence in the multimorbid, and the number of co-occurring diseases for each disease, were calculated. We used LCA to explore multimorbidity patterns. The optimal number of latent classes was determined based on the lowest Consistent Akaike Information Criterion (CAIC) and the adjusted Bayesian-Schwarz Information Criterion (adjusted BIC) [35-37], clinical significance, and interpretability [38]. After selection of an optimal model, each respondent was assigned to the class for which s/he had the highest computed membership probability. An average posterior probability greater than 70% indicates an optimal fit [39]. The characteristics of respondents in different latent classes were compared using the chi-squared test for categorical variables and ANOVA for continuous variables. In addition, multinomial logistic regression was performed to assess the association between each sociodemographic factor (age, gender, household income, educational level, and occupation) and latent class membership. Associations were assessed using odds ratios (ORs) with 95% confidence intervals (CIs). Each association was adjusted in terms of the other variables in the multivariate analysis. Regression analysis (using SAS PROC SURVEYREG) was performed to assess whether the health-related quality of life varied by latent class membership. The least square mean of the EQ-5D index score (with the 95% CI) was estimated for each latent class. The adjusted model considered gender, age, household income, educational level, and occupation. Bonferroni post-hoc testing was performed after all pairwise comparisons. As the EQ-5D index score is not normally distributed, it was log-transformed prior to analysis and then back-transformed. All tests were two-tailed, and a p-value <0.05 was regarded as statistically significant. All statistical analyses were performed with the aid of SAS software (version 9.4; SAS Institute, Cary, NC, USA). All estimates were subjected to sample weighting to reflect the complexity of the KNHANES sampling design.

### Ethics

The KNHANES was approved by the institutional review board of the KCDCP (approval nos. 2013-07CON-03-4C, 2013-12EXP-03-5C, and 2015-01-02-6C). Written informed consent was obtained from all participants.

## Results

A total of 8,370 participants aged over 50 years were included in analysis. The mean age was 62.5 years, and 46% were male. Of all respondents, 39% had two or more chronic diseases; the mean number of chronic diseases in multimorbid subjects was 2.6. Table 1 lists the prevalence and the proportions of multimorbidity, and the average number of comorbid diseases for each of the 10 chronic diseases included in the analysis. Hypertension (36.4%), dyslipidemia (22.1%), osteoarthritis (19.8%), and diabetes mellitus (14.4%) were the most prevalent diseases and at least one of the 10 diseases existed as a multimorbidity in over 60% of multimorbid patients; stroke occurred in 87.1% (the highest) and tuberculosis in 63.2% (the lowest). The number of co-occurring diseases varied between 2.2 and 2.9 depending on the index disease. Table 2 summarizes the LCA model fits. When up to four latent classes were considered, the smallest adjusted BIC (two-class model: 929.18; three-class: 797.28; four-class: 801.36) and CAIC (two-class model: 1016.91; three-class: 930.97; four-class: 981.01) were those of the three-class model; these classes were *relatively healthy*, those with *cardiometabolic conditions*, and those with *arthritis, asthma, allergic rhinitis, depression, and thyroid disease* as revealed by the estimated probabilities of any particular chronic disease given membership of a latent class. Every respondent was assigned to one of the three classes based on the highest membership probability. The *relatively healthy* group included those with a low prevalence of all evaluated chronic conditions. The dyslipidemia response probabilities were very similar for the two multimorbid groups but dyslipidemia was classified as a *cardiometabolic condition* based on the clinical nature of the disease; thus, the *cardiometabolic conditions* group was populated by those with high probabilities of hypertension, dyslipidemia, stroke, and diabetes mellitus. The class membership probabilities were 60.4, 27.8, and 11.8% respectively, and the average posterior probabilities for all three classes exceeded 70% (Table 3). The mean age of the *cardiometabolic conditions* group was the highest (67 years) but the mean number of co-occurring chronic conditions was highest (2.9) in the *arthritis, asthma, allergy, depression, and thyroid disease* group. Both the *cardiometabolic conditions* group (40.2% male vs. 59.8% female) and the *arthritis, asthma, allergy, depression, and thyroid disease* group had higher proportions of females (17.7 vs. 82.3%); males constituted over 50% of the *relatively healthy* group (51.4 vs. 48.6%). In particular, most subjects were female in the *arthritis, asthma, allergy, depression, and thyroid disease* group (82.3%). Socioeconomic status was similar in the *cardiometabolic conditions* group and the *arthritis, asthma, allergy, depression, and thyroid disease* group; both groups evidenced higher proportions (compared to the overall figures) of individuals with lower household incomes, lower levels of education, and of unemployed status (Table 4).

**Table 1.** The Prevalence and Characteristics of Certain Chronic Diseases in 8,370 KNHANES 2013-2015 Respondents Aged 50 Years and Older.

| Disease | Prevalence | Cases with multimorbidty | Number of co-occurring diseases |
| --- | --- | --- | --- |
|  | % (SE) | % | Mean (SE) |
| Hypertension | 36.4 (0.39) | 69.4 | 2.2 (0.02) |
| Dyslipidemia | 22.1 (0.26) | 83.2 | 2.6 (0.03) |
| Stroke | 4.1 (0.11) | 87.1 | 2.9 (0.07) |
| Osteoarthritis | 19.8 (0.26) | 73.9 | 2.4 (0.03) |
| Tuberculosis | 5.9 (0.14) | 63.2 | 2.2 (0.06) |
| Asthma | 3.6 (0.09) | 83.5 | 2.8 (0.08) |
| Allergic rhinitis | 8.1 (0.12) | 67.6 | 2.3 (0.06) |
| Depression | 5.6 (0.14) | 74.7 | 2.6 (0.07) |
| Diabetes mellitus | 14.4 (0.18) | 84.4 | 2.7 (0.04) |
| Thyroid disease | 3.9 (0.10) | 74.9 | 2.5 (0.08) |

KNHANES, Korea National Health and Nutrition Examination Survey; SE, Standard error

**Table 2.**
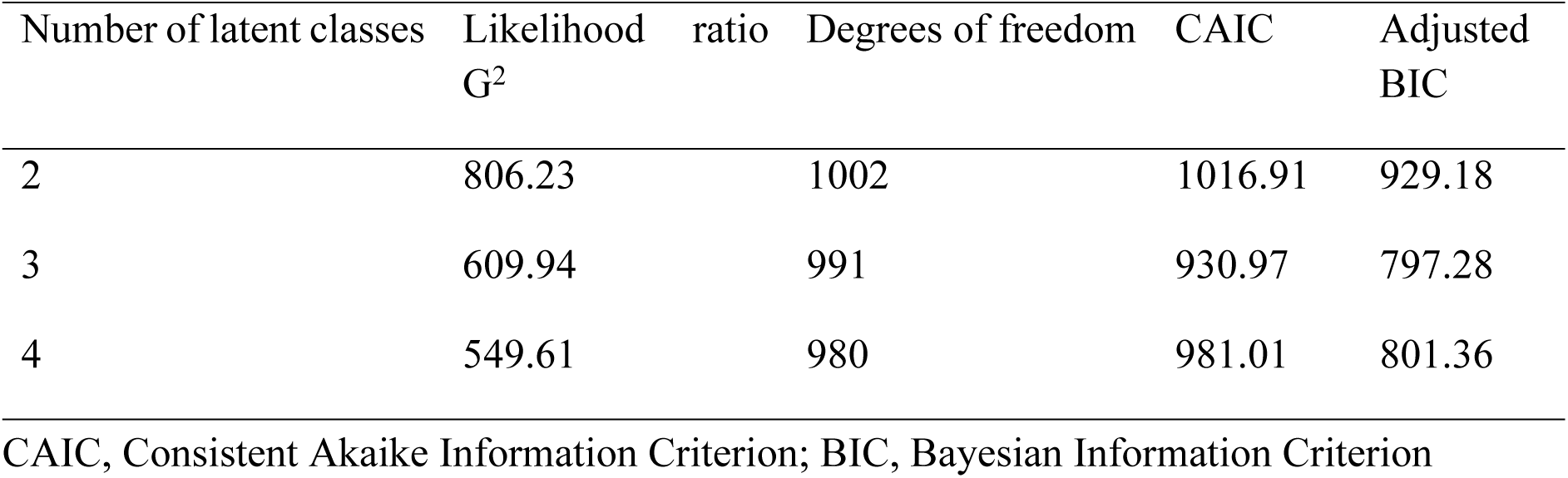
A Comparison of the Fit Statistics of Models Featuring Latent Class Analyses.

**Table 3.** Class Membership and Item Response Probabilities of the Three Latent Classes.

| Characteristics | Latent Class |  |  |
| --- | --- | --- | --- |
|  | Relatively<br>healthy | Cardiometabolic<br>conditions | Arthritis, asthma,<br>allergy, depression,<br>thyroid |
| Class membership probabilities (%) | 60.4 | 27.8 | 11.8 |
| Item response probabilities (%) |  |  |  |
| Hypertension | 0.18 | 0.75 | 0.42 |
| Dyslipidemia | 0.06 | 0.45 | 0.49 |
| Stroke | 0.01 | 0.12 | 0.02 |
| Osteoarthritis | 0.12 | 0.26 | 0.47 |
| Tuberculosis | 0.06 | 0.07 | 0.05 |
| Asthma | 0.02 | 0.04 | 0.14 |
| Allergic rhinitis | 0.07 | 0.04 | 0.24 |
| Depression | 0.03 | 0.04 | 0.21 |
| Diabetes mellitus | 0.04 | 0.39 | 0.13 |
| Thyroid disease | 0.02 | 0.03 | 0.14 |

**Table 4.**
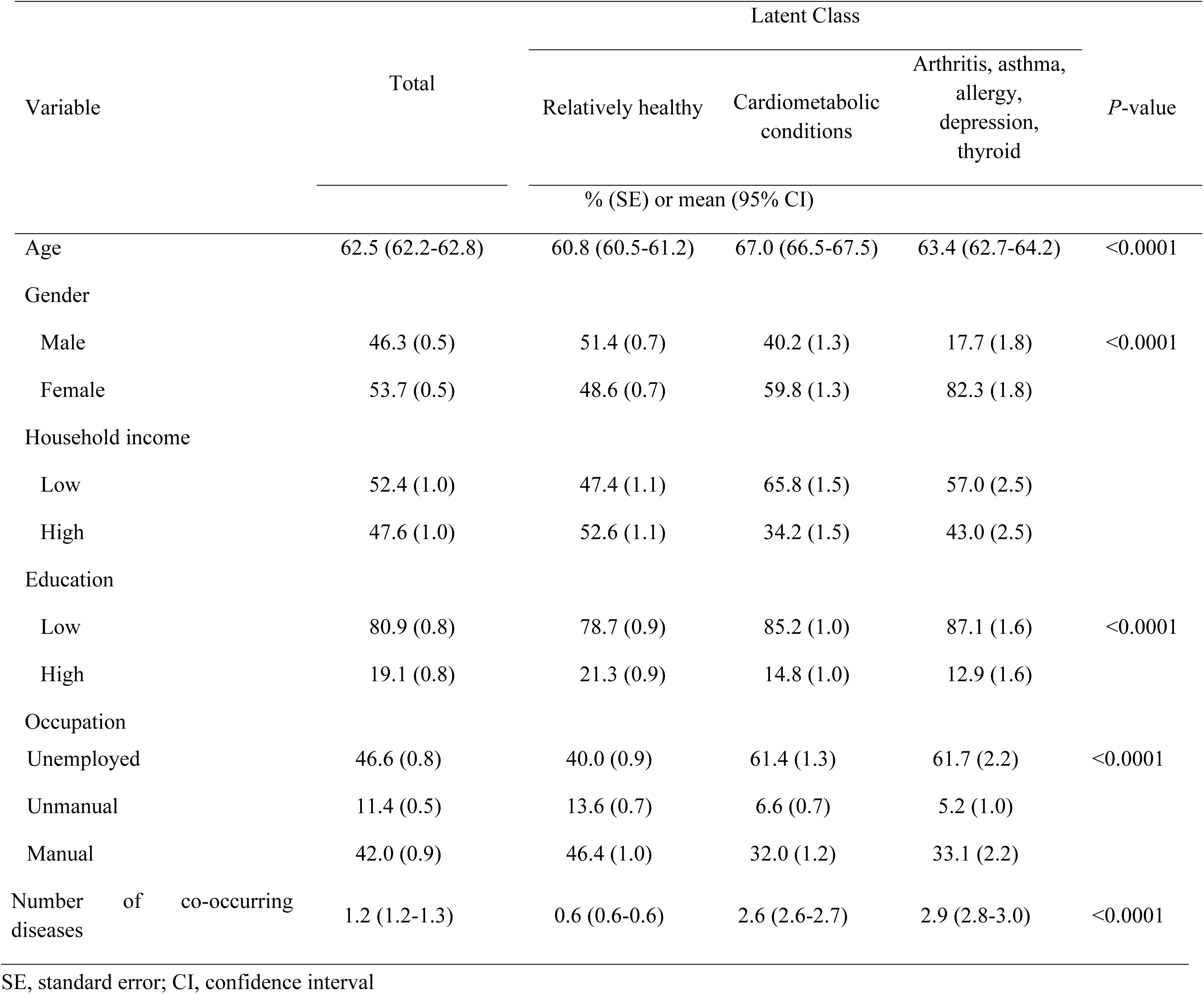
Characteristics of Study Respondents by Latent Class Membership.

Multinomial logistic regression analysis adjusted for age, gender, educational level, household income, and occupation showed that age increased the risk of being in either of the multimorbid groups compared to the *relatively healthy* group. The risks of membership of the *cardiometabolic conditions* group (OR=1.32, 95% CI=1.15-1.51) and the *arthritis, asthma, allergy, depression, and thyroid disease* group (OR=4.32, 95% CI=3.30-5.66) were significantly higher for females than males. Lower household income (OR=1.32, 95% CI=1.14-1.52) and lower educational level (OR=1.25, 95% CI=1.05-1.49) significantly increased the risk of membership of the *cardiometabolic conditions* group. In addition, unemployed status increased the risk of membership of both the *cardiometabolic conditions* group and the *arthritis, asthma, allergy, depression, and thyroid disease* group (Table 5).

**Table 5.** Sociodemographic Factors of the Latent Classes as Revealed by Multinomial Logistic Regression.

| Variable | Latent Class |  |  |
| --- | --- | --- | --- |
|  | Relatively healthy | Cardiometabolic conditions | Arthritis, asthma, allergy, depression, thyroid |
| Odds Ratio (95% Confidence interval) |  |  |  |
| **Unadjusted** |  |  |  |
| Age | 1 | 1.07 (1.07-1.08) | 1.03 (1.02-1.04) |
| Gender (ref=Male) |  |  |  |
| Female | 1 | 1.57 (1.39-1.78) | 4.90 (3.80-6.32) |
| Household Income (ref=High) |  |  |  |
| Low | 1 | 2.13 (1.88-2.42) | 1.47 (1.21-1.80) |
| Education (ref=High) |  |  |  |
| Low | 1 | 1.55 (1.32-1.82) | 1.83 (1.40-2.39) |
| Occupation (ref=Unemployed) |  |  |  |
| Unmanual | 1 | 0.32 (0.25-0.41) | 0.25 (0.16-0.38) |
| Manual | 1 | 0.45 (0.40-0.51) | 0.46 (0.38-0.57) |
| **Adjusted** |  |  |  |
| Age | 1 | 1.06 (1.05-1.07) | 1.02 (1.01-1.03) |
| Gender (ref=Male) |  |  |  |
| Female | 1 | 1.32 (1.15-1.51) | 4.32 (3.30-5.66) |
| Household Income (ref=High) |  |  |  |
| Low | 1 | 1.32 (1.14-1.52) | 1.10 (0.88-1.37) |
| Education (ref=High) |  |  |  |
| Low | 1 | 1.25 (1.05-1.49) | 1.12 (0.85-1.49) |
| Occupation (ref=Unemployed) |  |  |  |
| Unmanual | 1 | 0.80 (0.61-1.06) | 0.56 (0.36-0.86) |
| Manual | 1 | 0.70 (0.61-0.81) | 0.68 (0.54-0.85) |

CI, confidence interval

Table 6 shows the association between class membership and health-related quality-of-life. When the least-square means of the health-related quality-of-life of the latent classes were compared after adjustment for age, gender, household income, educational level, and occupation, individuals in the *arthritis, asthma, allergy, depression, and thyroid disease* group evidenced a significantly lower health-related quality-of-life (0.84) than the other two groups (0.93 for the *relatively healthy* group; 0.88 for the *cardiometabolic conditions* group) as revealed by Bonferroni post-hoc analysis.

**Table 6.** Associations between Multimorbidity Patterns and EQ-5D Index Scores.

|  | Latent Class |  |  | *P*-value<sup>a</sup> |
| --- | --- | --- | --- | --- |
|  | Relatively healthy | Cardiometabolic conditions | Arthritis, asthma, allergy, depression, thyroid |  |
| Unadjusted | 0.92 (0.91, 0.92) | 0.84 (0.83, 0.84) | 0.80 (0.79, 0.82) | <0.0001 |
| Adjusted |  |  |  |  |
| Model 1 | 0.91 (0.90, 0.92) | 0.86 (0.85, 0.87) | 0.82 (0.81, 0.84) | <0.0001 |
| Model 2 | 0.93 (0.92, 0.94) | 0.88 (0.87, 0.89) | 0.84 (0.83, 0.86) | <0.0001 |

EQ-5D, European Quality of Life 5 Dimension
Values are presented as geometric mean (95% confidence interval)
<sup>a</sup>*P*-value was calculated from Bonferroni post-hoc testing
Model 1: Age, gender
Model 2: Age, gender, household income, education, occupation

## Discussion

We used LCA to identify three distinct multimorbidity patterns in the general Korean population: (1) a relatively healthy group; (2) a group with cardiocerebrovascular conditions including hypertension, dyslipidemia, stroke, and diabetes mellitus; and, (3) a group with arthritis, asthma, allergic rhinitis, depression, and thyroid disease. We found that individuals of the three classes exhibited different sociodemographic characteristics and varied in terms of health-related quality of life. It is difficult to directly compare our results to those of previous studies given the methodological differences in terms of study setting, disease spectrum and number, demographic factors, baseline health status of participants, and the statistical methods used [29], but the multimorbidity patterns we identified are in general agreement with those of prior works. Such similarities may indicate that chronic diseases aggregate because they share underlying risk factors (so one disease increases the risk of another) or that causalities are shared [19]. We found that hypertension, dyslipidemia, stroke, and diabetes mellitus were very likely to co-occur, as have several prior studies evaluating subjects with (predominantly) cardiocerebrovascular conditions [17,19,21,40-42]. It is known that hypertension, dyslipidemia, and diabetes mellitus are risk factors for cerebrovascular diseases, and may precede such diseases [43-46]. Although the mechanism behind the combination of arthritis, asthma, allergic rhinitis, depression, and thyroid disease remains unclear, similar groupings were evident in other studies. Arthritis and depression were co-grouped by Simoes et al. [19], asthma and allergy by Larsen et al. [47], and arthritis, respiratory diseases, and psychiatric symptoms formed a latent class in the work of Islam [17,42].

Most previous studies on multimorbidity used a non-model approach such as counts of the most common disease combinations and their observed-to-expected ratios [48-50]; such methods are too simplistic. Recently, several studies have used statistical approaches such as cluster analysis [31,51,52], factor analysis [21,25,40,41], and LCA [17,22,36,42,53-55] to identify nonrandom multimorbidity clusters. We used LCA to identify subgroups based on structural equation modeling [56]. LCA is better than conventional clustering because LCA employs probability-based classification methods to choose an optimal number of classes based on various diagnostic tests [27,57]. This allowed us to group individuals into a limited number of latent classes and then analyze the differences between the classes. Our analysis not only indicated what other diseases were likely to co-occur in subjects with certain diseases but also allowed us to explore whether individuals with certain sociodemographic characteristics were more vulnerable to multimorbidity and the associated adverse health outcomes. The multimorbid groups were of lower socioeconomic status than the relatively healthy group. Individuals with cardiometabolic conditions were older than individuals in the *relatively healthy* and the *arthritis, asthma, allergy, depression, and thyroid disease* groups. In addition, a remarkable female predominance was evident in the *arthritis, asthma, allergy, depression, and thyroid disease* group. Female predominance in a musculoskeletal class and a headache-mental illness class was noted by Larsen et al. [47].

Furthermore, we found that older age, being female, and lower socioeconomic status increased the risk of membership of the *cardiometabolic conditions* group and/or the *arthritis, asthma, allergy, depression, and thyroid disease* group, as compared to the relatively healthy group. In particular, being female increased the risk of membership of the *arthritis, asthma, allergy, depression, and thyroid disease* group more than four-fold; also, individuals assigned to this latent class had a lower quality-of-life and were younger than members of other groups. Females are thus significantly more likely to develop the disease cluster of arthritis, asthma, allergy, depression, and thyroid disease than males and, thus, endure a lower quality-of-life for longer than males. Such gender inequality in healthy life expectancy, caused by multimorbidity, warrants public health interventions targeting the female elderly.

Our study had certain limitations. Analysis was based on a limited number of chronic diseases recorded by the KNHANES only from non-institutionalized Koreans who were able to participate in the survey. In addition, we only considered disease occurrence and, thus, not duration or severity. Finally, we cannot discuss causal relationships between diseases because the survey was cross-sectional in nature. However, using a large and nationally representative sample, we identified multimorbidity patterns, and associations between such patterns and both sociodemographic factors and health-related quality-of-life. Our findings deepen our understanding of non-random associations between diseases; this will aid the design of useful, effective, holistic healthcare and preventative strategies addressing the needs of multimorbid individuals and those at high risk of multimorbidity. In addition, given that the multimorbidity patterns are associated with poor quality-of-life and sociodemographic inequalities, targeted multimorbidity management is important to reduce the burden on the vulnerable population and to address the associated social inequalities.

